# Novel Diazirine Photoprobes for the Identification of Vancomycin-Binding Proteins

**DOI:** 10.1101/2023.06.15.545140

**Authors:** Photis Rotsides, Paula J. Lee, Nakoa Webber, Kimberly C. Grasty, Joris Beld, Patrick J. Loll

## Abstract

Vancomycin’s interactions with cellular targets drive its antimicrobial activity, and also trigger expression of resistance against the antibiotic. Interaction partners for vancomycin have previously been identified using photoaffinity probes, which have proven to be useful tools for exploring vancomycin’s interactome. This work seeks to develop diazirine-based vancomycin photoprobes that display enhanced specificity and bear fewer chemical modifications, as compared to previous photoprobes. Using proteins fused to vancomycin’s main cell-wall target, D-alanyl-D-alanine, we use mass spectrometry to show that these photoprobes specifically label known vancomycin-binding partners within minutes. In a complementary approach, we developed a Western-blot strategy targeting the vancomycin adduct of the photoprobes, eliminating the need for affinity tags and simplifying the analysis of photolabeling reactions. Together, the probes and identification strategy provide a novel and streamlined pipeline for identifying novel vancomycin-binding proteins.

**For Table of Contents Only.**

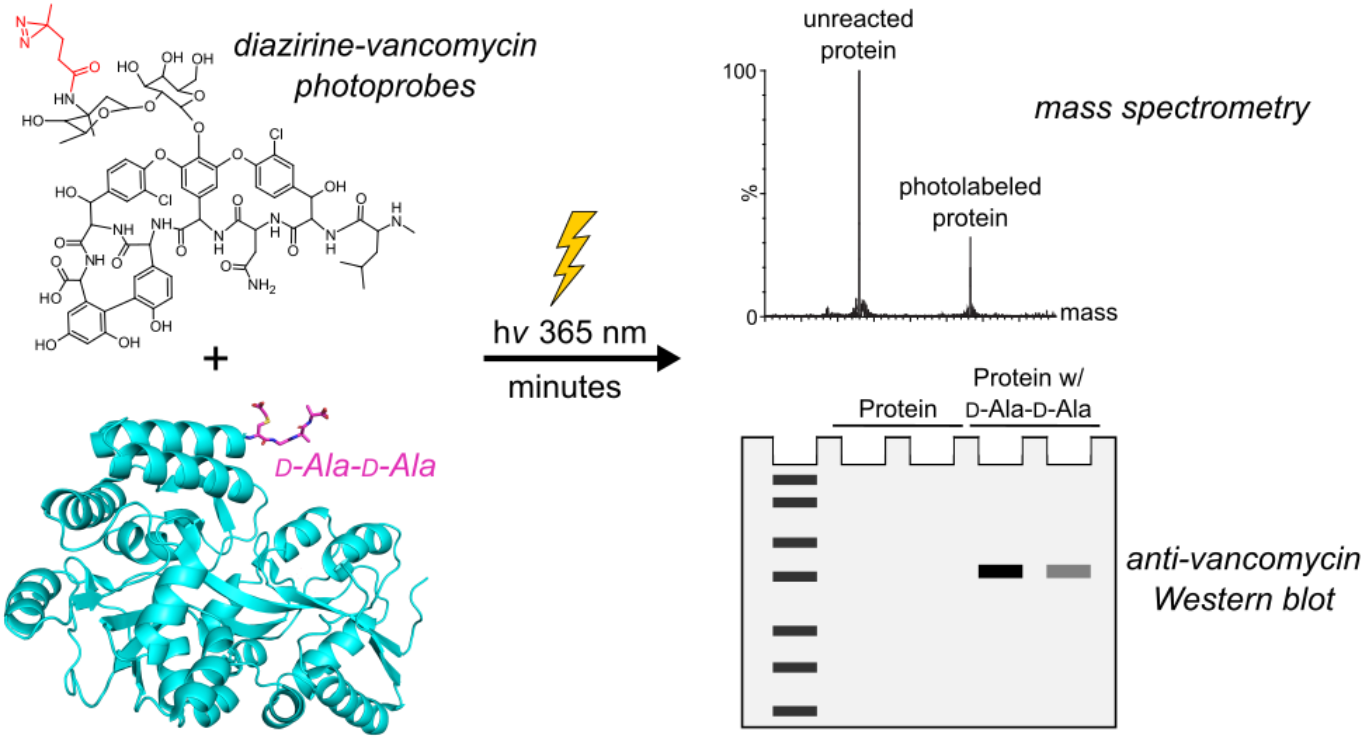

## Introduction

Vancomycin is a glycopeptide antibiotic that is used in healthcare settings across the globe.^1^ It has historically served as an antibiotic of last resort to treat persistent infections caused by Gram-positive microbes.^2-4^ However, decades of clinical use have led to the emergence of vancomycin resistance in various human pathogens, most notably <u>v</u>ancomycin-resistant *<u>E</u>nterococci* (VRE).^5-8^ VRE infections cause thousands of deaths each year, and lead to extended hospital stays and increased treatment costs. Accordingly, VRE have been identified as high-priority pathogens for the development of new therapeutic approaches.^9-12^Efforts to develop novel VRE-targeted therapies will benefit from a thorough understanding of all of vancomycin’s interaction partners. Vancomycin binds the D-alanyl-D-alanine (D-Ala-D-Ala) moiety of the cell-wall precursor Lipid II, thereby interfering with peptidoglycan synthesis.^13-16^ D-Ala-D-Ala is the substrate for the transpeptidase enzyme, and by sequestering this peptide, vancomycin inhibits the cross-linking reaction required to form a rigid cell wall, leaving the bacteria susceptible to lysis.^17^ VRE elude vancomycin’s effects by remodeling D-Ala-D-Ala to D-alanyl-D-lactate or D-alanyl-D-serine, neither of which is efficiently recognized by the antibiotic.^18-25^ The VRE resistance phenotype is regulated by VanS, a membrane-bound sensor histidine kinase.^26^ VanS detects the presence of the antibiotic and responds by initiating a signaling pathway that culminates in the expression of cell-wall remodeling enzymes.^26-28^ However, it remains unclear whether VanS proteins sense vancomycin by binding directly to the antibiotic, or through more indirect mechanisms.^6, 29, 30^ Complicating matters, nine different types of VRE have been described and the corresponding VanS proteins vary significantly in sequence, raising the possibility that some VanS proteins may bind directly to vancomycin, while others do not.^6, 31^ Furthermore, the vancomycin-protein interactome may extend beyond the VanS proteins, as vancomycin has been suggested to interact with additional proteins, including an autolysin and an ABC transporter.^32^ Thus, new tools that can aid in identifying and characterizing vancomycin’s molecular partners would be welcome.

One tool that has found wide application in profiling interacting partners is photoaffinity labeling.^33^ This approach has already been applied to vancomycin, having been used to probe the interaction between the antibiotic and the VanS protein from *Streptomyces coelicolor*, and for proteomic studies aimed at identifying novel interaction partners.^32, 34^ In both of these examples, the vancomycin photoprobes contained benzophenone groups as the photoactive species. Benzophenones are relatively bulky, and thus can potentially cause steric interference with interacting partners; they also require long periods of UV irradiation for effective protein labeling, which can damage the target molecule and increases the possibility of nonspecifically labeling bystander proteins.^33^ We therefore chose to explore diazirine-containing photoprobes. Diazirines are smaller and more flexible than benzophenones, decreasing the risk of steric clashes in protein-binding interfaces; they also require less UV exposure for activation, reducing the radiation burden on the sample.^35, 36^ Upon UV irradiation, diazirines form highly reactive and unstable carbenes, which last for only picoseconds before being quenched by water.^37-39^ These short half-lives ensure that the carbenes are extinguished before they can diffuse and react with off-target proteins, providing improved specificity over benzophenones.

Previously used vancomycin photoprobes also included biotin or alkyne handles to facilitate isolation of the photolabeled species. Such handles are useful when isolating targets from complex cellular mixtures, but contribute additional bulk that can disrupt interactions. Encouragingly, in the examples cited above these modifications did not impair vancomycin’s antibiotic activity, nor did they alter the antibiotic’s ability to bind to D-Ala-D-Ala *in vitro*; however, it is still possible that they might interfere with other biologically relevant interactions. We therefore questioned whether it would be possible to avoid incorporating additional tags such as biotin into the photoprobe. Instead, we chose to rely upon direct immunodetection of vancomycin, using commercially available antibodies. Together, these strategies have allowed us to produce novel and effective vancomycin photoprobes, which improve our ability to identify vancomycin-interacting proteins and complement the existing repertoire of tools.

## Results and Discussion

### Development and identification of diazirine-containing vancomycin photoprobes

Targeting vancomycin’s amine groups has proven to be a straightforward synthetic strategy for introducing additional groups onto the antibiotic.^40^ Vancomycin contains a primary amine on its vancosamine sugar and a secondary amine at its methylated N-terminus (Figure 1A); modifications at both positions are known to preserve antibiotic activity, making these plausible sites for introducing the photolabel. We used a diazirine derivative containing an NHS ester to acylate the antibiotic, producing multiple species that could be separated using reverse-phase high-pressure liquid chromatography (RP-HPLC) (Figure 1B). Mass spectrometry (MS) revealed that two of the isolated compounds had masses consistent with singly-labeled species, presumably reflecting attachment of the diazirine at either the vancosamine or the N-terminus (compounds **2** and **3**, respectively; Figure 1C). An additional species was obtained, having a mass consistent with doubly-labeled product (**4**), which was not pursued further.

**Figure 1.**
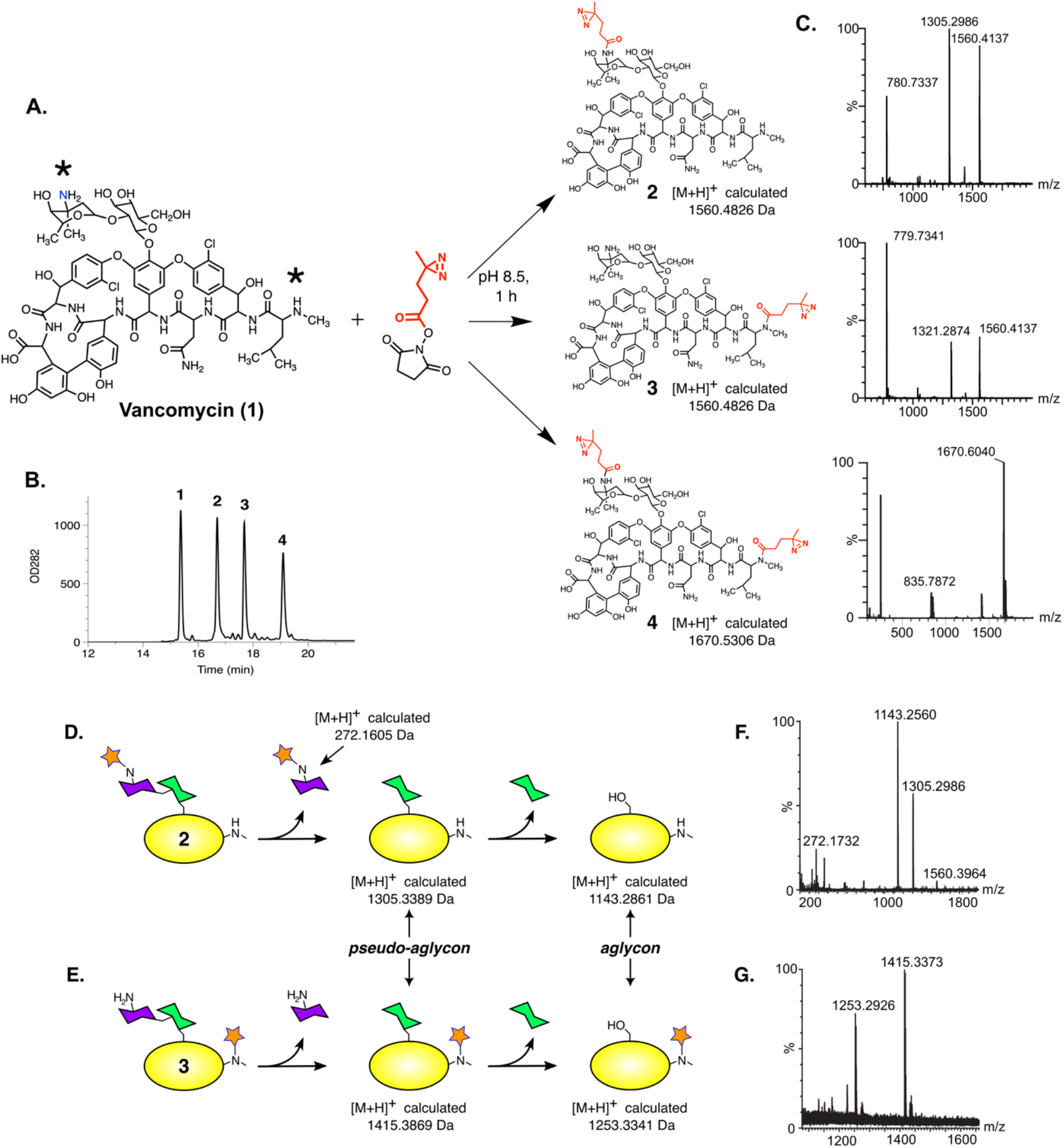
Preparation of vancomycin-diazirine photolabeling reagents. (*A*) Reaction of vancomycin with a diazirine-NHS ester yields three products. The NHS ester is expected to react at two positions (indicated by asterisks): A primary amine on the vancosamine sugar, and the secondary amine at vancomycin’s N-terminus. (*B*) The products and starting material were separated using RP-HPLC. (*C*) Mass spectrometry was used to confirm the presence of the adduct in the three products. For compound **3**, in addition to the singly-and doubly-charged versions of the molecule, an additional peak is seen at 1321.2874, which likely represents a breakdown product of the labeled pseudo-aglycon (see Figure S2). (*D & E*) Schematic drawings showing the anticipated acid hydrolysis products and their predicted masses for photoprobe **2**, in which the diazirine group is attached to the vancosamine sugar (panel *D*), or photoprobe **3**, with the diazirine attached to the N-terminus (panel *E*). The diazirine is represented by the orange star. (*F & G)* Mass spectra of the acid-hydrolysis products.

We next sought to determine the locations of the diazirine group in the two singly-labeled species. One of the species provided a clue about its identity in its native mass spectrum, which showed a peak for the pseudo-aglycon lacking the diazirine adduct, suggesting it was compound **2** (Figure 1C; see Figure S1 for a definition of the pseudo-aglycon and aglycon. Formation of the pseudo-aglycon frequently occurs during ionization of vancomycin^41^). Conversely, the other singly-labeled species contained a peak at 1321.2874 in its native mass spectrum, consistent with a degradation product derived from the pseudo-aglycon of compound **3** (Figure S2). The MS/MS data were consistent with these assignments (Figure S3).

To provide independent confirmation of the identities of the two singly-labeled species, we used acid hydrolysis to remove one or both of the antibiotic’s sugar molecules (Figure 1D & 1E; Figure S1). In the case of **2**, removal of the vancosamine generates pseudo-aglycon and aglycon species lacking the diazirine, whereas for **3** both species retain the diazirine. Compound **2** was readily identified, as it gave the unlabeled pseudo-aglycon and aglycon species, along with a molecule having the appropriate mass for a diazirine-labeled vancosamine sugar (Figure 1F). Likewise, hydrolysis of the other singly-labeled sample yielded pseudo-aglycon and aglycon species containing the diazirine group, confirming its identity as **3** (Figure 1G).

### Diazirine-vancomycin photoprobes specifically label D-Ala-D-Ala-containing proteins

To test whether our photoprobes could label known vancomycin-binding proteins, we created constructs in which D-Ala-D-Ala peptides were fused to the C-termini of different proteins that served as carriers. Specifically, we used native chemical ligation to couple an L-Cys-L-Lys-D-Ala-D-Ala peptide to C-terminal thioesters of maltose-binding protein (MBP), T4 lysozyme (T4L), and ubiquitin (Ub).^42^ Following coupling, the cysteine was alkylated with iodoacetic acid. The resulting protein-peptide conjugates contain a C-terminal sequence that closely mimics the peptide moiety of Lipid II. All three of these protein-peptide conjugates bind vancomycin with low-micromolar affinities, and they have been used to determine crystal structures of vancomycin and other glycopeptide antibiotics bound to their peptide targets.^42, 43^ Notably, in the absence of the D-Ala-D-Ala-containing peptide, none of the three protein carriers themselves bind to vancomycin.

For photolabeling reactions, proteins were mixed with either photoprobe **2** or **3**, and then irradiated with 365-nm UV light, after which the products were analyzed by MS. As expected, no photolabeling was observed using control proteins lacking the D-Ala-D-Ala peptide (Figure S4). All of the D-Ala-D-Ala-containing constructs were photolabeled, albeit with varying efficiencies. Substantial labeling of MBP-D-Ala-D-Ala (MBP-DADA) was observed with **2**, whereas labeling by **3** was only roughly half as efficient (Figure 2A). **2** was similarly more efficient than **3** in labeling T4L-D-Ala-D-Ala (T4L-DADA) and Ub-D-Ala-D-Ala (Ub-DADA; Figure 2B-C); overall, labeling was most efficient for MBP-DADA.

**Figure 2.**
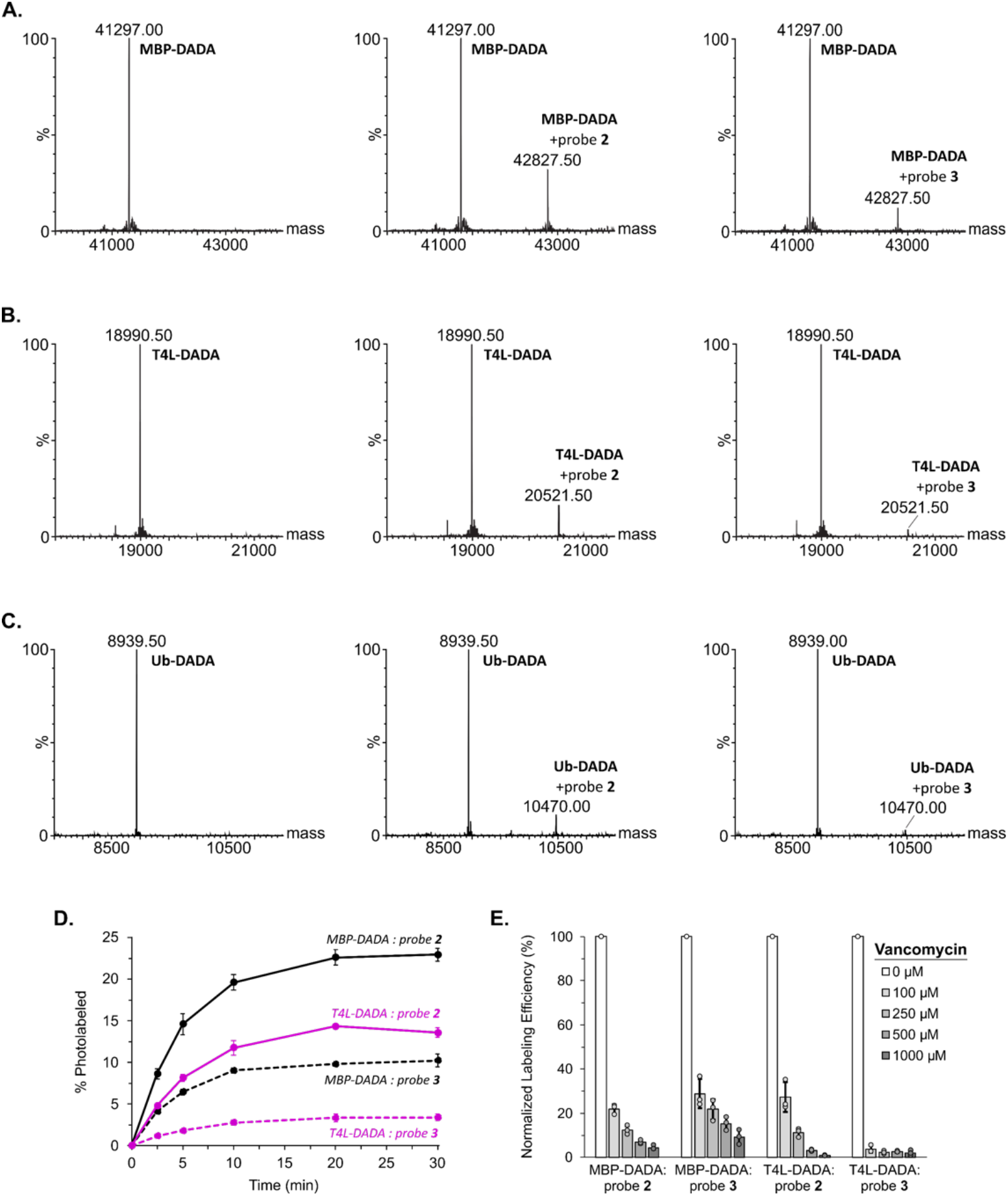
Photolabeling of protein-peptide conjugates containing D-Ala-D-Ala. (A) MBP-DADA (predicted mass 41297.91), (B) T4L-DADA (predicted mass 18990.75), and (C) Ub-DADA (predicted mass 8939.26) were mixed with buffer (left), photoprobe **2** (middle), or photoprobe **3** (right) and irradiated with UV light for 30 minutes. (D) Time-course of photolabeling reactions for MBP-DADA and T4L-DADA. The percentage of photolabeled protein was determined by the ratio of the unlabeled and photolabeled peak intensities in the mass spectra. Error bars represent one standard deviation (*n* = 3). (E) Competition experiments with vancomycin. Titration series for MBP-DADA and T4L-DADA were created containing either photoprobe **2** or **3** with increasing amounts of vancomycin. Samples were irradiated with UV light for four minutes, after which the efficiency of photolabeling was determined by mass spectrometry. Efficiency was normalized to the value observed in the absence of vancomycin. Error bars represent one standard deviation (*n* = 3).

We also observed varying photolabeling efficiencies within a given sample. For example, one of our MBP-DADA preparations was incompletely alkylated by iodoacetic acid, resulting in a sample containing roughly equal amounts of the S-carboxymethyl-modified cysteine and the unmodified cysteine (Figure S5). When this preparation was labeled with probe **2**, the strong majority of the labeled species corresponding to the unmodified cysteine, implying that photolabel **2** bound preferentially to peptides containing an unalkylated cysteine. In contrast, probe **3** showed no preference, with the photolabeled species being split equally between proteins containing modified and unmodified cysteines.

Kinetic experiments with the MBP-DADA and T4L-DADA constructs indicated that under these conditions, photolabeling reached a maximum by 20 minutes, with significant levels of labeling being observed after only 2.5 minutes (Figure 2D). This is significantly shorter than the two-hour exposures required for benzophenone-containing vancomycin photoprobes, which should presumably translate into an enhanced specificity for the new diazirine probes. To test the specificity of labeling, we asked whether unlabeled vancomycin could compete with the photoprobes. For both compounds **2** and **3**, addition of unlabeled vancomycin reduced photolabeling efficiency in a dose-dependent manner for both MBP-DADA and T4L-DADA (Figure 2E), suggesting that both photoprobes bind specifically to the vancomycin-binding site. Overall, these results demonstrate that the diazirine photoprobes are able to efficiently and specifically label known vancomycin-binding partners. They also reveal that photolabeling efficiency depends on the location of the photoreactive group, highlighting the utility having multiple probes in which the diazirine is attached at different positions.

### Anti-vancomycin Western blot for detecting photolabeled proteins

We anticipate that some vancomycin-binding partners, like VanS, will be membrane-bound. Membrane-protein preparations typically include lipids and/or detergents, which complicate mass spectrometric analysis. Therefore, we sought to develop an alternate technique for detecting photolabeled proteins. Because our photoprobes introduce a covalent vancomycin adduct, we reasoned that an anti-vancomycin antibody could be used to identify labeled proteins via standard immunodetection methods. Direct detection of the vancomycin molecule would have the additional advantage of abolishing the need for other tags (e.g., biotin or click handles), thereby reducing the size and complexity of the photoprobe. For these reasons, we used a commercial anti-vancomycin antibody to develop a novel Western-blot strategy for identifying photoprobe-labeled proteins.

Proteins were subjected to photolabeling, after which they were analyzed by denaturing SDS-PAGE and Western blotting. The labeling patterns observed in the Western blots agreed remarkably well with those seen using MS. Thus, all three D-Ala-D-Ala conjugates were seen to be labeled by **2** and **3** (Figure 3A), whereas proteins lacking the D-Ala-D-Ala group were not labeled by either compound. Labeling by probe **2** was much more efficient than probe 3 (Figure S6), consistent with the weaker intensities observed for probe **3**-labeled peaks in the MS data.

**Figure 3.**
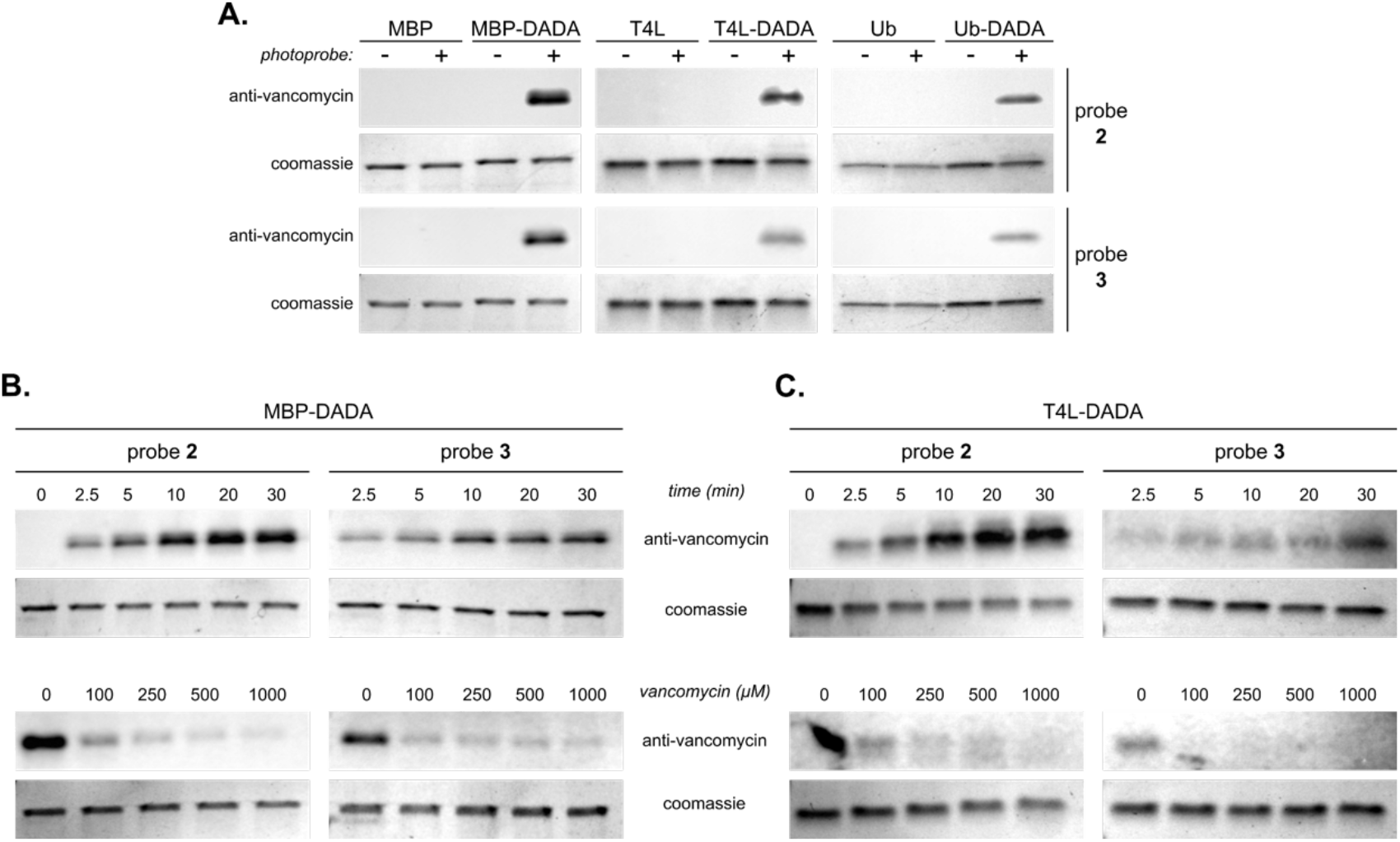
Photolabel Western blot using an anti-vancomycin antibody. (A) Western blots of photolabeling reactions for DADA-containing proteins, as well as control proteins lacking DADA. (B&C) Western blots showing time courses of photolabeling (upper panels) and competition experiments with vancomycin (lower panels) for MBP-DADA (panel B) and T4L-DADA (panel C). Coomassie-stained gels are shown as loading controls.

Kinetic and competition profiles obtained by Western blotting were also consistent with those observed using MS (Figure 3B & 3C). For both photoprobes, labeling of MBP-DADA and T4L-DADA was detectable within 2.5 minutes of UV exposure, and reached a plateau around 20 minutes. In both cases, a dose-dependent reduction in the Western-blot signal was seen when increasing amounts of free vancomycin were added to the photolabeling reactions, reinforcing that the signal is due to specific binding of the probes. Hence, this new Western-blotting approach provides a simple and powerful way to identify proteins labeled by vancomycin photoprobes and is likely to prove especially advantageous when studying vancomycin interactions with membrane proteins.

### Structural analysis of photoprobe interactions with D-Ala-D-Ala proteins

We wondered if the differences in labeling efficiency observed for the different probes and different protein-peptide conjugates could be rationalized in structural terms. To explore this question, we constructed structural models for photoprobes **2** and **3** bound to each of the three protein-peptide conjugates used. A crystal structure is already available for T4L-DADA bound to vancomycin (PDB ID 3RUN);^42^ for MBP-DADA and Ub-DADA, we used crystal structures of the proteins bound to teicoplanin and dalbavancin, respectively, as starting points for the modeling (PDB IDs 3VJF & 3RUL).^42, 43^

In the case of MBP-DADA, the primary amine on the vancosamine sugar lies close to MBP’s C-terminal alpha-helix, explaining why a diazirine attached at this position (probe **2**) robustly labels this protein (Figure 4A). On the other hand, the antibiotic’s N-terminus is oriented away from the body of MBP and projects outward into solvent; hence a diazirine attached at this position (probe **3**) will be farther from the protein, consistent with the lower efficiency of labeling associated with this probe-protein pair. The structure also offers an explanation for the preferential labeling of protein with an unalkylated cysteine by probe **2**: The S-carboxymethyl group on the modified cysteine lies immediately adjacent to the vancosamine sugar, and thus may interfere sterically with the diazirine group; in contrast, the smaller unmodified cysteine side chain offers less steric hindrance, suggesting **2** will bind preferentially to the unalkylated protein.

**Figure 4.**
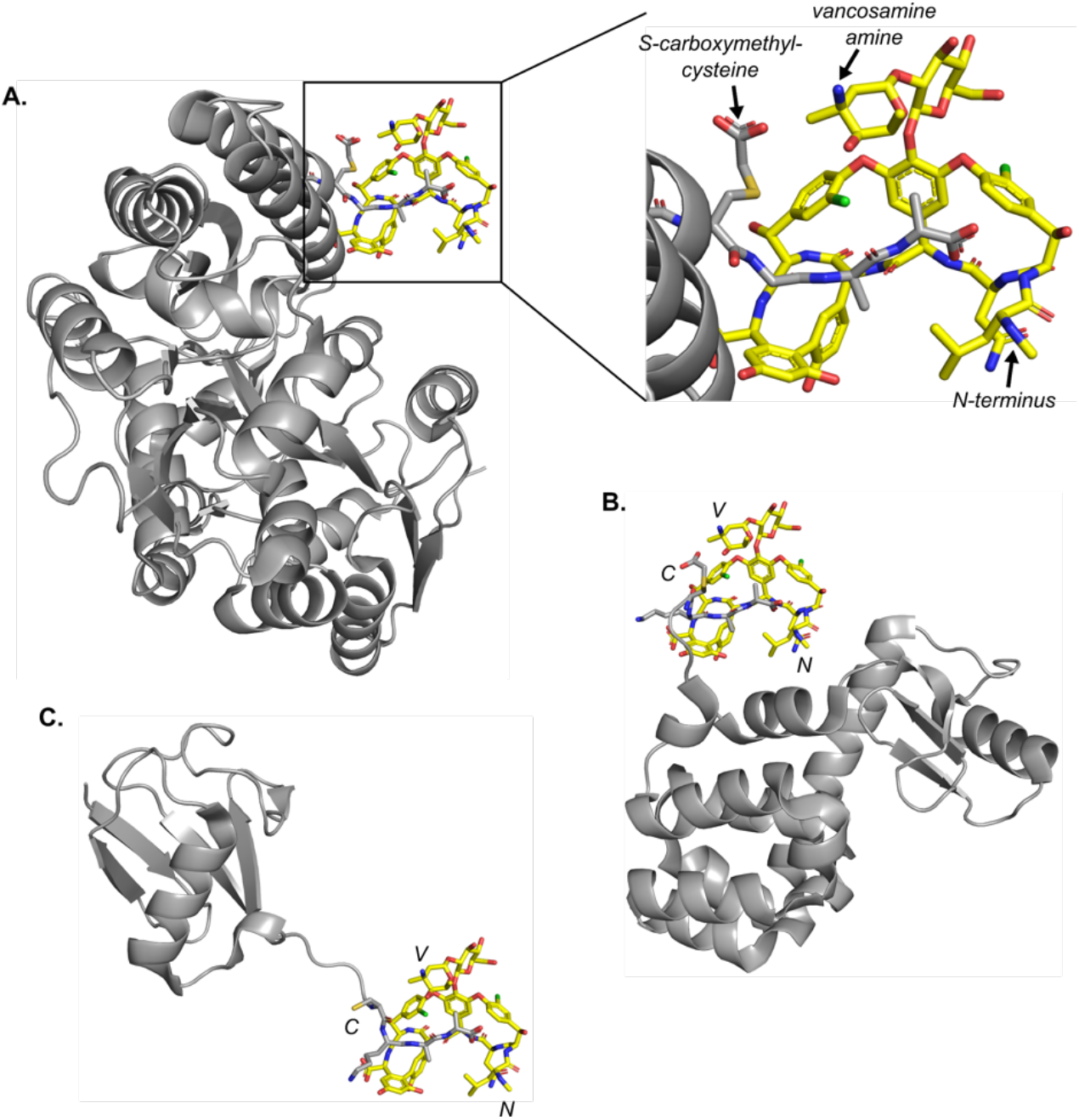
Structural modeling of photoprobe interactions with protein-peptide conjugates. (A) Model of the complex of vancomycin bound to MBP-DADA (left). The inset shows a close-up view of the interaction region, highlighting relative positions of the two diazirine attachment sites and showing the modified cysteine residue in the protein-peptide conjugate. (B) Crystal structure of T4L-DADA bound to vancomycin (PDB ID: 3RUN). (C) Model of the complex of Ub-DADA bound to vancomycin. In all panels, vancomycin is shown in yellow sticks, while the protein-peptide conjugates are colored gray, with the fused Cys-Lys-D-Ala-D-Ala peptides being shown as sticks. The positions of the vancosamine amine and N-terminus of vancomycin are indicated by V and N, respectively, while the cysteine of the ligated peptide is indicated by C.

In the crystal structure of T4L-DADA bound to vancomycin, vancomycin’s N-terminus is close to the protein’s first alpha helix, suggesting a potential steric clash with the diazirine of probe **3** (Figure 4B), possibly explaining its poor labeling efficiency for this protein. Finally, in the case of Ub-DADA, the structure reveals that the protein’s C-terminus extends outward, away from the body of the molecule; thus, when vancomycin binds to the D-Ala-D-Ala group, diazirine groups attached to either the antibiotic’s N-terminus or its vancosamine sugar will lie far from the core of the ubiquitin molecule, explaining why Ub-DADA was less efficiently labeled by both probes, as compared to the other D-Ala-D-Ala conjugates (Figure 4C).

The structural mode by which vancomycin recognizes D-Ala-D-Ala probably also contributes to the stronger labeling of the protein-peptide conjugates by probe **2** versus probe **3**. When D-Ala-D-Ala is bound to vancomycin, its C-terminus lies next to the antibiotic’s N-terminal amine. By definition, the C-terminal residues are the ones farthest away from the remainder of the protein; this likely biases probe **3** toward positions that are more removed from the bulk of the protein than is true for probe **2**. Overall, then, our structural knowledge of how vancomycin binds to the protein-peptide conjugates is consistent with the observed photolabeling patterns. Importantly, the structural context underscores the importance of close spatial proximity between the photoreactive group and the protein target, as well as good structural complementarity in the probe-target pair.

In conclusion, we report novel diazirine-vancomycin photoprobes that can specifically label known vancomycin-binding proteins, while requiring only a fraction of the UV burden needed for previous probes; ultimately, this should reduce sample damage and decrease the chance of labeling off-target proteins. We have prepared two complementary photoprobes having reactive groups located at different positions, increasing the likelihood that at least one probe will be able to engage a given target without steric clashes. Chances of such clashes are further minimized by small size of the diazirine groups and by our antibody strategy for recognizing the vancomycin adduct, which allowed us to generate probes having minimal modifications to the vancomycin molecule. Going forward, this antibody-based approach may also prove useful for proteomic studies, enabling the isolation of photolabeled proteins without the need for an affinity tag.

## Supporting Information

Detailed methods for photoprobe synthesis and characterization, protein purification and preparation of protein-peptide conjugates, photolabeling, mass spectrometry, and Western-blot assays.

## Acknowledgements

This work was supported by grant 1R01AI148679 from the National Institute of Allergy and Infectious Disease, National Institutes of Health.

