## Supplemental Information for "Novel Diazirine Photoprobes for the Identification of Vancomycin-Binding Proteins"

### *List of Contents*

***Detailed methods***

Photoprobe preparation & characterization ........ S2

LC-MS methods ........ S2

Protein expression & purification ........ S2

Native chemical ligation ........ S4

Photolabeling reactions ........ S4

Western-blot detection of photolabeling ........ S4

***Fig. S1.*** Formation of pseudo-aglycon and aglycon by acid hydrolysis ........ S5

***Fig. S2.*** Degradation scheme for pseudo-aglycon of **3**. ........ S6

***Fig. S3.*** MS/MS degradation products of **2** and **3**. ........ S7

***Fig. S4.*** Photolabeling data for negative-control proteins ........ S8

***Fig. S5.*** Effect of cysteine alkylation on MBP-DADA photolabeling ........ S9

***Fig. S6.*** Western blot comparing MBP-DADA photolabeling efficiency with **2** and **3** ........ S10

***References*** ........ S11

### Methods

*Preparation of vancomycin photoprobes*. The following solutions were prepared: 1 M dibasic sodium phosphate, pH 8.5; 10 mg/mL vancomycin in water; and 10% (w/v) succinimidyl 4,4'-azipentanoate (Thermo Scientific) in DMSO. To 50 µL of the vancomycin solution (500 µg, 0.35 µmol) was added 5 µL of the phosphate buffer and 3.5 µL of the diazirine solution (1.07 µmol, 3.1 equivalents). The mixture was incubated at room temperature, protected from light, for one hour, after which 5 µL of 1 M hydroxylamine was added. The incubation was continued for an additional hour and then the reaction stopped by freezing at -20°C. Upon thawing, the reaction mixture was filtered through a 0.45 µm filter and purified by reverse-phase chromatography on a C18 column (Ultrasphere 5 ODS, 1.0 x 25 cm, Hichrom Ltd.), using a gradient from 5-100% acetonitrile in a mobile phase containing 0.25% formic acid.

*Characterization of photoprobe position using acid hydrolysis*. Hydrolysis reactions were prepared as described with slight modification.[1] Briefly, 20 µL of 1 mg/mL solutions of each photoprobe were mixed with 3.5 µL of 4N HCl then boiled for 3 minutes. The samples were then cooled on ice for 5 minutes, after which 1 mL of 5 mM ammonium bicarbonate pH 8.0 was added to each sample to neutralize the acid. The masses of the hydrolysis products were determined by MS to identify the location of the diazirine group for each photoprobe.

*Liquid-chromatography mass spectrometry*. Molecules were analyzed on a Waters Acquity I-Class UPLC system coupled to a Synapt G2Si HDMS mass spectrometer in positive ion mode with a heated electrospray ionization (ESI) source in a Z-spray configuration. For proteins, LC separation was performed on a Waters Acquity UPLC Protein BEH C4 1.7 µm 2.1 x 50 mm column maintained at 80 °C. After an initial 1 min at 0.4 ml/min of 95/5 A/B, a 0.2 ml/min gradient to 5/95 A/B in 3.5 min was used to separate proteins, followed by washing and reconditioning the column. For vancomycin species, LC separation was performed on a Waters Acquity UPLC BEH C18 1.7 µm 2.1 x 50 mm column maintained at 40 °C using an 0.6 ml/min gradient of 95/5 to 15/85 A/B over the course of four minutes. Eluent A is 0.1% v/v formic acid in water and B is 0.1% v/v formic acid in acetonitrile. Conditions on the mass spectrometer were as follows: capillary voltage 0.5 kV, sampling cone 40 V, source offset 80 V, source 120 °C, desolvation 250 °C, cone gas 0 L/h, desolvation gas 1000 L/h and nebulizer 6.5 bar. The analyzer was operated in resolution mode and low energy data was collected between 100 and 2000 Da at 0.2 sec scan time. For vancomycin derivatives, MSe data was collected using a 20-40V ramp trap collision energy. Masses were extracted from the TOF MS TICs using a 0.005 Da abs width. Protein ESI data was deconvoluted using MaxEnt1 in Masslynx 4.1 (Waters Corporation).

*Expression and purification of protein-intein fusions.* Preparation of the histidine-tagged intein vectors used in this work was previously described.[2] The MBP and ubiquitin constructs were transformed into BL21(DE3) cells (New England BioLabs) and the T4L construct was transformed into BL21(DE3) pLysS cells (Novagen). Cells were grown in ZYP-5052 auto-inducing media[3] with 100 µg/mL ampicillin, with additional 34 µg/mL chloramphenicol for the T4L construct. Proteins were expressed at 24 °C for 24 hours. Unless otherwise stated, all purification steps were carried out at 4 °C or on ice. Cells were harvested by centrifugation, washed in MilliQ water and stored at -80 °C. Pellets were resuspended in Buffer A (50 mM sodium phosphate, 250 mM NaCl, 10 mM imidazole, pH 7), lysed using an Emulsiflex C5 high-pressure homogenizer (Avestin, Inc., Ottawa, Canada), and centrifuged for 30 min at 9,000*g*. The supernatant was further clarified by ultracentrifugation for 50 min at 240,000*g*. The supernatant was passed over a HiTrap IMAC HP column (Cytiva #17092005) charged with nickel and equilibrated in Buffer A. The column was washed with 0.1% v/v Triton-X100 in Buffer A (10 bed volumes), re-equilibrated in Buffer A, and eluted with Buffer B (25 mM sodium acetate pH 3.6, 500 mM NaCl). The fractions were neutralized with 1 M HEPES pH 8, and the most concentrated fractions were pooled. EDTA was added to a concentration of 2 mM and intein cleavage was initiated by adding dry 2-mercaptoethanesulfonate (MESNA) to a final concentration of 500 mM. The mixture was incubated overnight at room temperature and then dialyzed against Buffer C (25 mM MES pH 6.5, 250 mM NaCl; two changes, 3 hours each). The reaction mixture was passed over the HiTrap IMAC HP column, which had previously been equilibrated in Buffer C. The column flow-through was collected and concentrated for the native protein ligation to >60 mg/ml for the T4L and MBP and >25 mg/ml for ubiquitin.

*Native protein ligation*. Synthetic peptides were fused to protein carriers using native protein ligation as described with minor alterations.[4] Briefly, the freshly MESNA-cleaved, concentrated protein was incubated for two days at room temperature in 0.1 M HEPES pH 8, 500 mM NaCl, 500 mM MESNA, and 2- to 10-fold molar excess of synthetic L-Cys-L-Lys-D-Ala-D-Ala (Biomatik). In the case of the MBP and T4L constructs, following the ligation, 10 mM TCEP was added to the ligation mixture to fully reduce the protein. Reducing agents were then removed by two rounds of desalting on a HiPrep 26/10 desalting column (GE Healthcare) equilibrated in alkylation buffer (0.1M sodium phosphate pH 8, 150 mM NaCl, 5 mM EDTA), after which a 10-fold molar excess of iodoacetic acid was added to the protein and incubated for two hours at room temperature. Finally, the protein-peptide fusions were dialyzed against 20 mM HEPES pH 7, 25 mM NaCl and concentrated. For the ubiquitin construct, after the ligation the sample was dialyzed into sodium acetate pH 4.6, 5 mM DTT and loaded onto a HiTrap SP-HP anion exchange column (GE Lifesciences) equilibrated in the same buffer. The protein was eluted with a linear gradient from 0-30% of elution buffer (sodium acetate pH 4.6, 5 mM DTT, 1M NaCl). The second peak to elute, corresponding to the ubiquitin-peptide fusion, was dialyzed twice against alkylation buffer for 1 hour at room temperature, then a 10-fold molar excess of iodoacetic acid was added. The solution was incubated for two hours at room temperature, dialyzed against sodium acetate pH 4.6, 5 mM DTT, and then run as previously on the HiTrap SP-HP anion exchange column. Fractions containing the ubiquitin-peptide fusion were dialyzed against 20 mM HEPES pH 7, 25 mM NaCl and concentrated. Masses of the final, purified protein-peptide fusions were verified by mass spectrometry.

*Photolabeling reactions*. 20-µL reactions containing 20 µM protein and 60 µM photoprobe were prepared in 20 mM Tris pH 7.5, 100 mM NaCl. The reactions were placed into glass depression spot plates, which were placed atop a heavy aluminum plate packed in ice, to keep reactions cold and to limit evaporation. Samples were irradiated at 365 nm using a Stratagene Stratalinker 2400 UV Crosslinker for the times specified. For competition experiments with vancomycin, samples were irradiated for four minutes. All samples were diluted to 0.01 mg/mL for subsequent analysis.

*Western-blot analysis*. 100 ng of protein were loaded onto a 12% SDS-PAGE gel and electrophoresed at 180 V. While the gel was running, a 0.2 µm PVDF membrane (Cytiva #10600021) was prepared by soaking in 100% methanol for one minute then in water for two minutes. The membrane was then equilibrated in transfer buffer for 15 minutes (25 mM Tris pH 8.3, 192 mM glycine, 15% methanol). Proteins were transferred from the gel to the membrane for one hour at 100 V, after which the membrane was rocked in blocking buffer (5% milk in 20 mM Tris pH 7.6, 150 mM NaCl, 0.1% Tween-20 (TBST)) for 30 minutes. The membrane was then rocked overnight at 4°C in a solution containing a sheep anti-vancomycin polyclonal antibody (Bio-Rad # 9520-0004) diluted 1:1000 in blocking buffer. The following day, the membrane was washed in TBST 3 x 10 min and then rocked at room temperature for one hour in solution containing a rabbit anti-sheep HRP-conjugated secondary antibody (Invitrogen # 31480), diluted 1:5000 in blocking buffer. The membrane was then washed in TBST 3 x 10 min and placed in peroxidase substrate solution (Pierce #32209) for 1 minute before chemiluminescent detection.

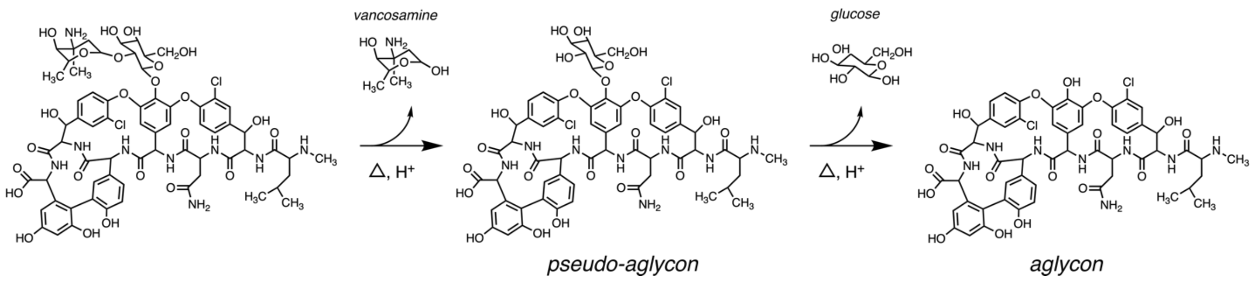


***Figure S1.*** Formation of the vancomycin pseudo-aglycon and aglycon by acid hydrolysis.

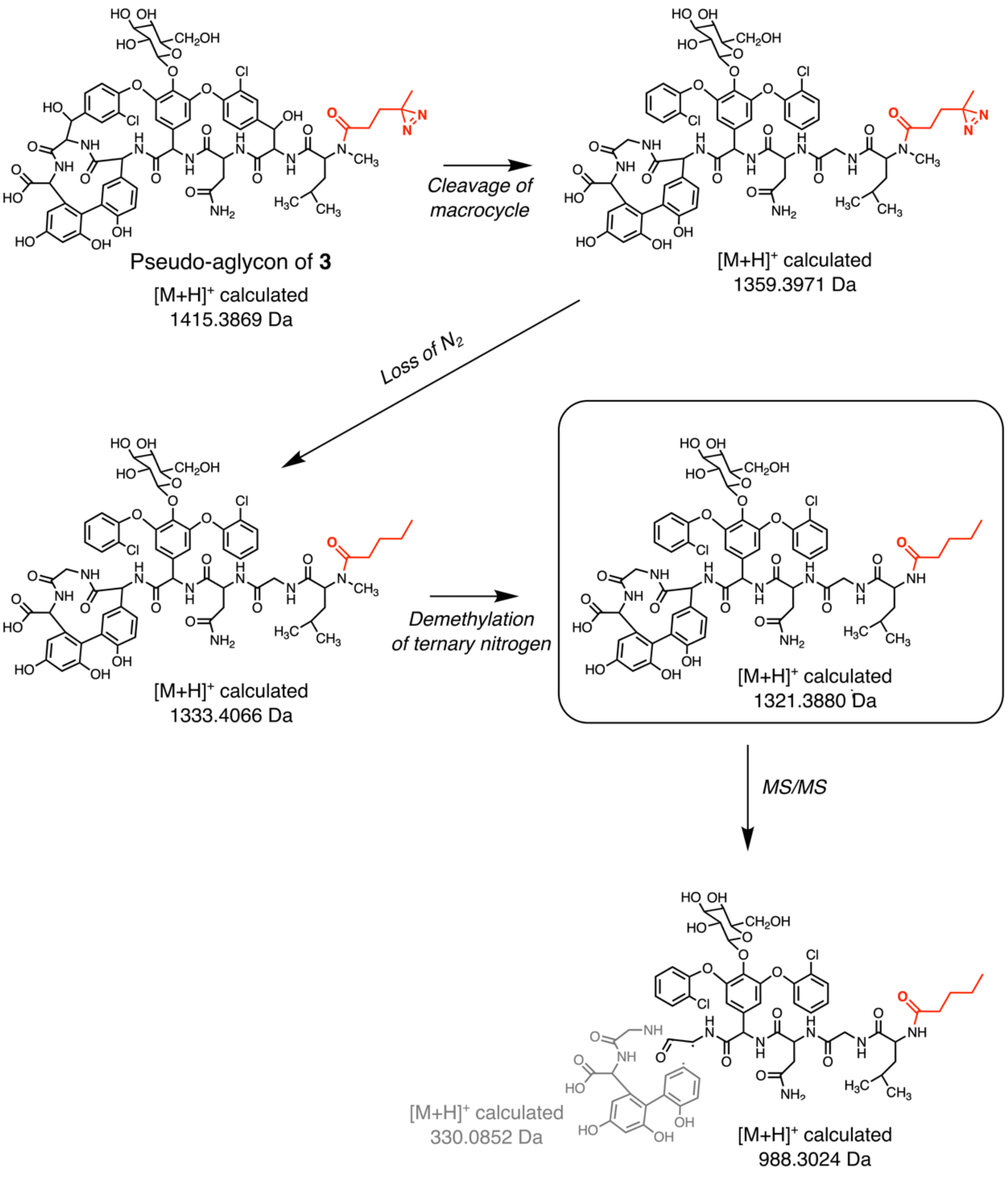


**Figure S2**. *Degradation scheme for pseudo-aglycon of **3***. The native mass spectrum of probe **3** shows a species with m/z = 1321.2874 (Figure 1C), produced by the scheme shown. Starting with the pseudo-aglycon, cleavage of the macrocycle, loss of $N_2$ from the diazirine, and loss of the methyl group from the ternary nitrogen yield a species with the appropriate mass. Supporting this conjecture, further cleavage in the MS/MS experiment is predicted to yield a species with mass 988.3024, and a strong peak at m/z = 988.1713 is observed in the MS/MS data (not shown).

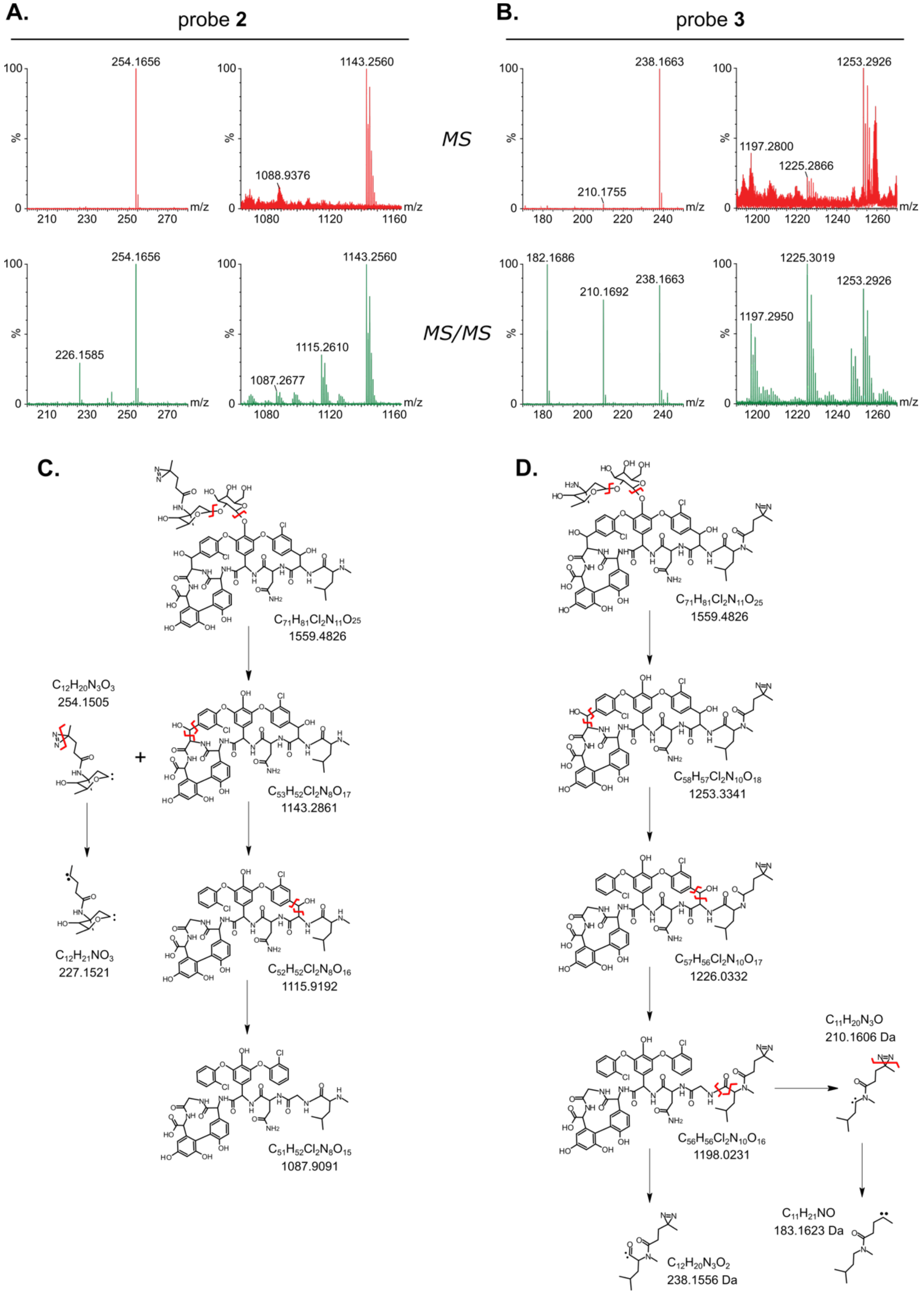


**Figure S3.** *MS/MS degradation products of **2** and **3**.* (A & B) Native (top) and tandem (bottom) mass spectra of **2** and **3**. (C & D) Degradation pathways for **2** (panel C) and **3** (panel D), highlighting the products observed in the mass spectra. Red lines represent fragmentation sites for subsequent degradation products.

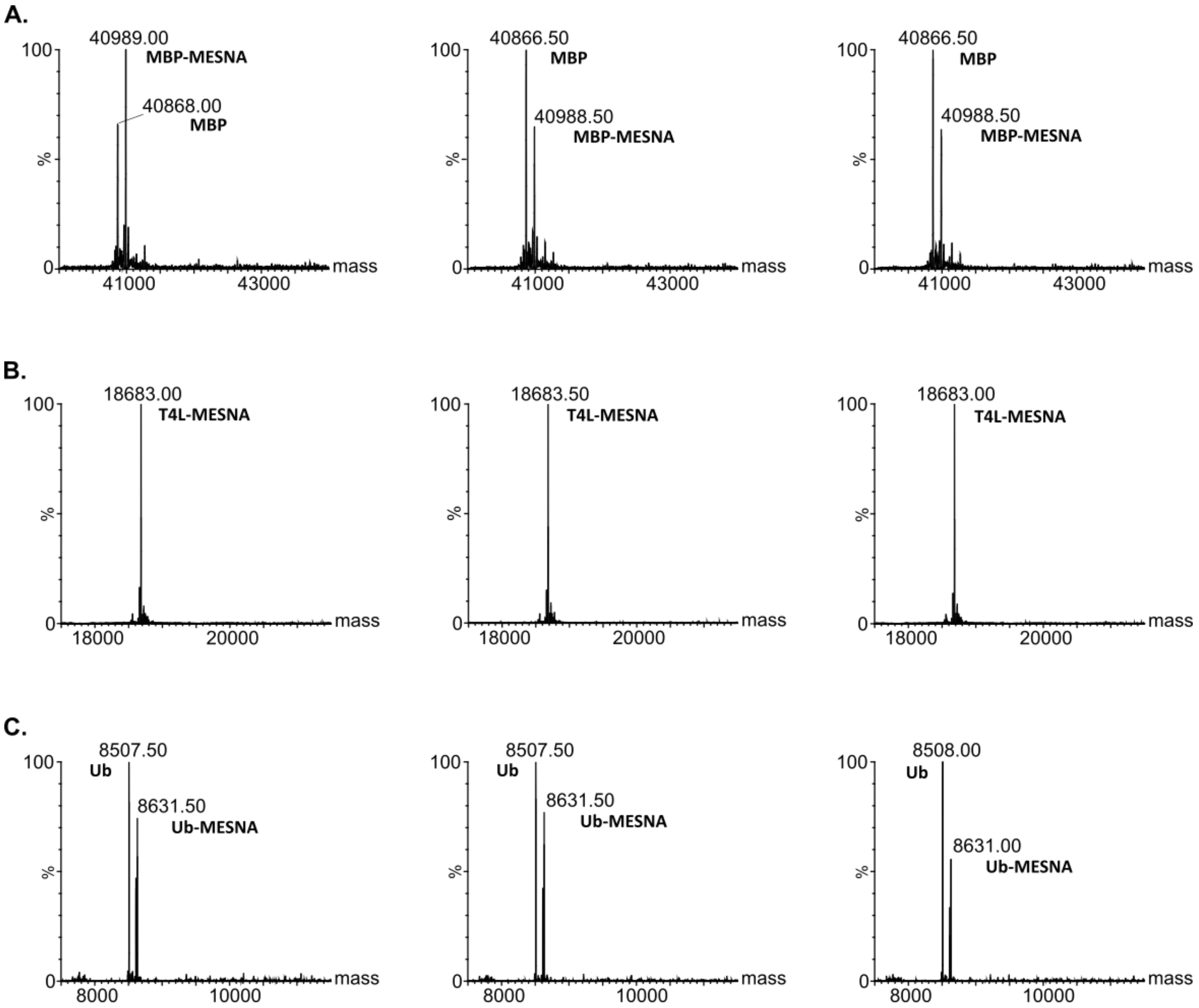


**Figure S4.** *Photolabeling data for negative-control proteins*. (A) MBP, (B) T4L, and (C) Ub were mixed with buffer (left), photoprobe **2** (middle), or photoprobe **3** (right) and irradiated with UV light for 30 minutes. MS was used to determine the presence of photolabeled protein. Doublets observed in the mass spectra represent proteins containing either a C-terminal MESNA thioester (produced during intein cleavage) or a free C-terminal carboxylate (produced by hydrolysis of the thioester).

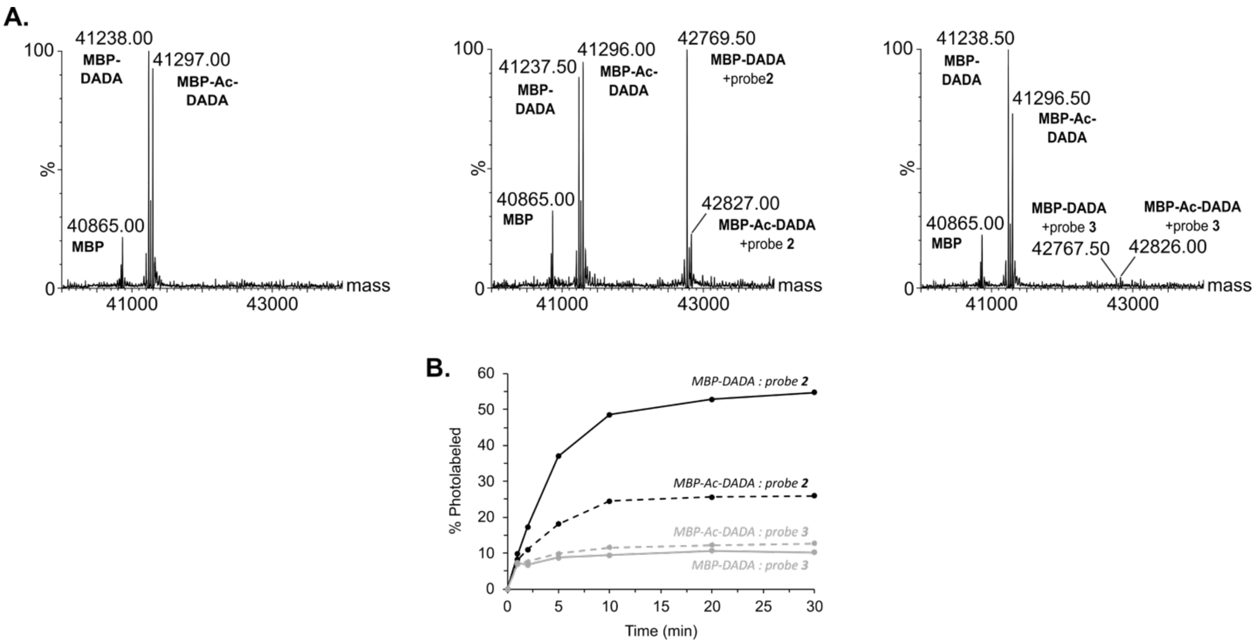


**Figure S5.** *Effect of cysteine alkylation on MBP-DADA photolabeling.* (A) Partially alkylated MBP-DADA was mixed with buffer (left), photoprobe **2** (middle), or photoprobe **3** (right) and irradiated with UV light for 30 minutes. MS was used to determine the presence of photolabeled protein. Doublets observed in the spectra represent species containing either an alkylated cysteine (MBP-Ac-DADA; expected mass 41297.91) or an unmodified cysteine (MBP-DADA; expected mass 41239.91). (B) Time-course of photolabel reactions. The percentage of photolabeled protein was determined by the ratio of the unlabeled and photolabeled peak intensities in the mass spectra.

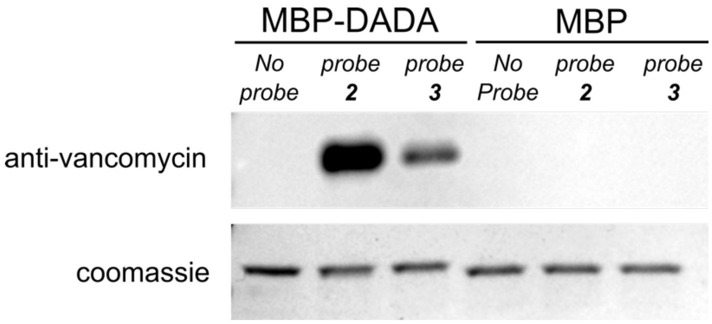


**Figure S6.** *Western blot directly comparing the efficiency of photolabeling for MBP-DADA and MBP, using probes **2** and **3**.* Proteins were photolabeled with either probe **2** or probe **3**. Equal quantities of photolabeled protein were loaded in each lane, and Western blotting was carried out as described above.
